# Serum protein profiling reveals hallmark-level aging trajectories and strain-specific resilience in CB6F1J and C57BL/6J male mice

**DOI:** 10.64898/2026.04.14.718493

**Authors:** Gerald Yu Liao, Christina Pettan-Brewer, Warren Ladiges

## Abstract

Aging is characterized by coordinated molecular and physiological changes across multiple biological systems, yet the ability to quantify these processes non-invasively within individuals remains limited. Here, we establish a framework for quantifying hallmark-level features of aging in mice using serum protein array profiles obtained from a single blood draw. Serum protein expression was profiled in groups of CB6F1J and C57BL/6J male mice at 8 and 32 months of age and mapped to established hallmarks of aging. Hallmark-level analyses revealed coordinated, pathway-specific changes in inflammatory, vascular, intercellular signaling, metabolic, and regenerative processes, with distinct patterns observed between strains. CB6F1J mice exhibited directional shifts across multiple pathways, while C57BL/6J mice showed broader but more heterogeneous changes. Cross-strain comparisons demonstrated shared pathway-level trends alongside variable protein-level concordance. This approach enables non-terminal assessment of aging within individuals and resolves heterogeneity in aging trajectories using minimally invasive sampling. These findings support the use of circulating protein signatures to quantify biological aging and provide a framework for translating non-invasive proteome-based assessment of aging to human studies.

## Introduction

Aging is the primary risk factor for most chronic diseases, yet the ability to quantify biological aging within individuals remains limited [1]. Foundational work has defined conserved molecular hallmarks of aging, including chronic inflammation, altered intercellular communication, and loss of proteostasis, that collectively drive functional decline [2–3]. In parallel, blood-based biomarkers such as epigenetic clocks and circulating proteomic signatures have demonstrate that systemic aging signals are measurable and correlate with chronological age and disease risk [4–5]. However, while these approaches demonstrate that aging-related signals are detectable in blood, they are typically optimized for predicting age or mortality risk rather than resolving underlying biological pathways or distinguishing resilience from decline. Moreover, current frameworks do not systematically map circulating biomarkers onto the canonical hallmarks of aging, limiting their interpretability and mechanistic relevance [6].

Murine models further underscore the complexity of aging biology, as strain-dependent differences produce divergent trajectories in lifespan, physiology, and molecular regulation. Hybrid strains such as CB6F1J exhibit distinct aging trajectories compared to inbred strains such as C57BL/6J, reflecting differences in genetic architecture and systemic regulation [7–9]. Despite these well-characterized discrepancies, it remains unclear whether such variation can be captured through non-invasive, repeatable measures capable of tracking biological aging over time [5].

Here, we present a framework for quantifying hallmark-level aging signatures using circulating protein array profiles obtained from a single blood draw and apply this approach to distinguish aging trajectories between CB6F1J and C57BL/6J male mice. This strategy enables repeated, non-terminal sampling within individuals, allowing direct comparison of biological states across the lifespan. Given the translational relevance of blood-based biomarkers in humans, this approach provides a potential foundation for assessing biological aging and resilience using minimally invasive methods.

We hypothesize that circulating protein array profiles can resolve hallmark-level features of aging and distinguish strain-specific aging trajectories, enabling non-invasive, quantification of biological aging within individuals.

## Methods

### Protein array profiling

Serum protein expression was quantified in groups of male CB6F1J and C57BL/6J mice, obtained from the National Institute on Aging’s Aged Rodent Colony, at 8 and 32 months of age using a multiplex antibody-based protein array (RayBiotech). Raw signal intensities were background-corrected and normalized to internal positive controls according to the manufacturer’s protocol. Proteins with signal intensities at or below background were excluded from downstream analyses.

Normalized expression values were log_2_-transformed following addition of a small pseudocount (0.01) to avoid undefined values. Age-associated changes were calculated as log_2_ fold change (log_2_FC) between 32-month and 8-month mice within each strain.

### Differential protein analysis

Differential expression between age groups was assessed for each protein using two-tailed unpaired t-tests within each strain. Proteins were classified as increased or decreased with age based on the sign of the log_2_FC. Statistical thresholds were denoted using conventional asterisk-based annotations (*, **, ***, ****). Analyses were performed across all detected proteins and separately within the subset meeting statistical thresholds.

### Functional annotation and hallmark mapping

Proteins were grouped into functional families based on canonical biological classifications, including cytokines, chemokines, extracellular matrix regulators, and angiogenic factors. These functional groups were mapped to established hallmarks of aging, including chronic inflammation, altered intercellular communication, dysregulated nutrient sensing, stem cell exhaustion, loss of proteostasis, cellular senescence, vascular dysfunction, and neurodegeneration, consistent with established frameworks [2–3]. Proteins with pleiotropic functions were assigned to multiple categories as appropriate.

### Hallmark-level analysis

Hallmark-level analyses were performed to quantify coordinated pathway-level changes. For each hallmark, the proportion of proteins exhibiting increased or decreased expression with age was calculated across all proteins and within the subset meeting statistical thresholds. Hallmark scores were defined as the mean log_2_FC across all proteins within each category.

### Cross-strain comparison

To assess shared and strain-specific aging signatures, protein-level and hallmark-level changes were compared between CB6F1J and C57BL/6J male mice. Concordance was quantified using Pearson correlation coefficients of log_2_FC values across strains, calculated for all proteins and for the subset meeting statistical thresholds.

### Statistical analysis

All analyses were performed in Microsoft Excel (Microsoft Corp.). Log_2_-transformed expression values were used for all downstream analyses. Pearson correlation coefficients were used to assess cross-strain concordance. Analyses restricted to statistically defined subsets were interpreted with consideration of reduced sample size. Figures were generated with R version 4.3.2 (R Foundation for Statistical Computing).

## Results

### CB6F1J male mice exhibited coordinated increases in inflammatory, vascular, and intercellular signaling proteins with concurrent reductions in metabolic and regenerative pathway components

In CB6F1J male mice, aging was associated with directional changes across multiple protein hallmarks (Table 1, Figures 1–3). Within inflammatory pathways, 52.7% (59/112) of proteins were increased with age, with 28.6% (32/112) contributing to the subset of most altered proteins. Hallmark scores were positive (log_2_FC = 0.48; significant proteins = 0.99).

**Table 1.** Distribution of directional and statistically defined protein changes across hallmarks of aging in CB6F1J male mice.

| Hallmark of Aging | Total N | ↑ Proteins (All) (N, %) | ↑ Proteins (Sig.) (N, %) | ↓ Proteins (All) (N, %) | ↓ Proteins (Sig.) (N, %) |
| --- | --- | --- | --- | --- | --- |
| Inflammaging | 112 | 59 (52.68) | 32 (28.57) | 52 (46.43) | 14 (12.50) |
| **Vascular Dysfunction** | 19 | 15 (78.95) | 10 (52.63) | 4 (21.05) | 2 (10.53) |
| **Altered Intracellular Communication** | 122 | 66 (54.10) | 33 (27.05) | 55 (45.08) | 14 (11.48) |
| **Stem Cell Exhaustion / Impaired Regeneration** | 44 | 20 (45.45) | 7 (15.91) | 24 (54.55) | 7 (15.91) |
| **Dysregulated Nutrient Sensing** | 26 | 11 (42.31) | 4 (15.38) | 15 (57.69) | 4 (15.38) |
| **Cellular Senescence** | 77 | 39 (50.65) | 21 (27.27) | 38 (49.35) | 11 (14.29) |
| **Loss of Proteostasis / Tissue Integrity** | 22 | 13 (59.09) | 3 (13.64) | 9 (40.91) | 4 (18.18) |
| **Neurodegeneration / Neural Dysfunction** | 5 | 3 (60.00) | 0 (0.00) | 2 (40.00) | 0 (0.00) |

**Figure 1.**
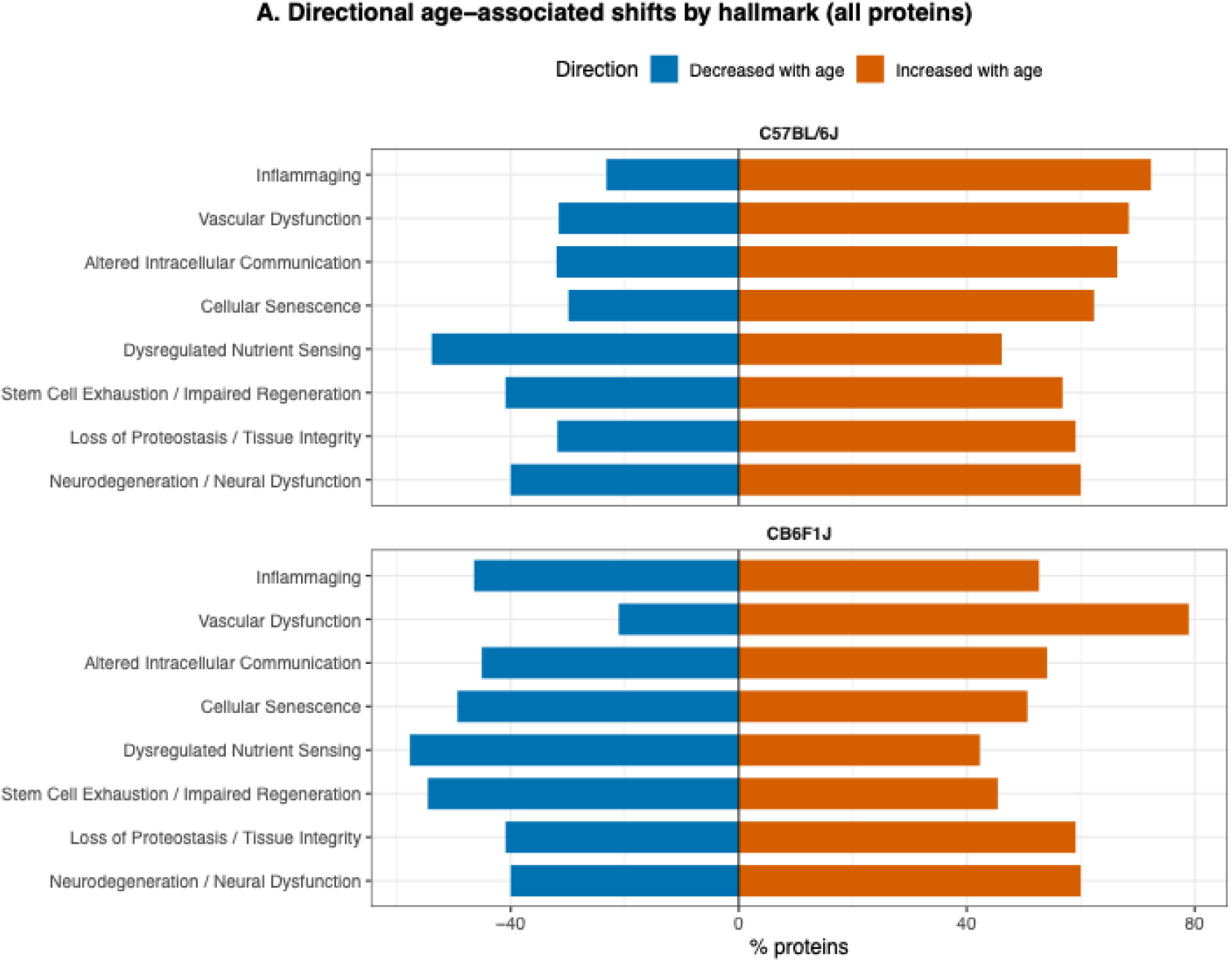
Directional age-associated shifts in circulating protein hallmarks across mouse strains for all analyzed proteins. For each hallmark of aging, the proportion of proteins exhibiting increased (orange) or decreased (blue) expression with age is shown for C57BL/6J (top) and CB6F1J (bottom) mice. Percentages were calculated across all detected proteins within each hallmark category. Positive values indicate the fraction of proteins increased with age, whereas negative values indicate the fraction decreased with age. Across both strains, inflammatory, vascular, and intercellular communication pathways show a predominance of increased proteins, while nutrient sensing and regenerative pathways display a higher proportion of decreased proteins. Differences in the relative distribution of directional changes are observed between strains across multiple hallmarks.

**Figure 2.**
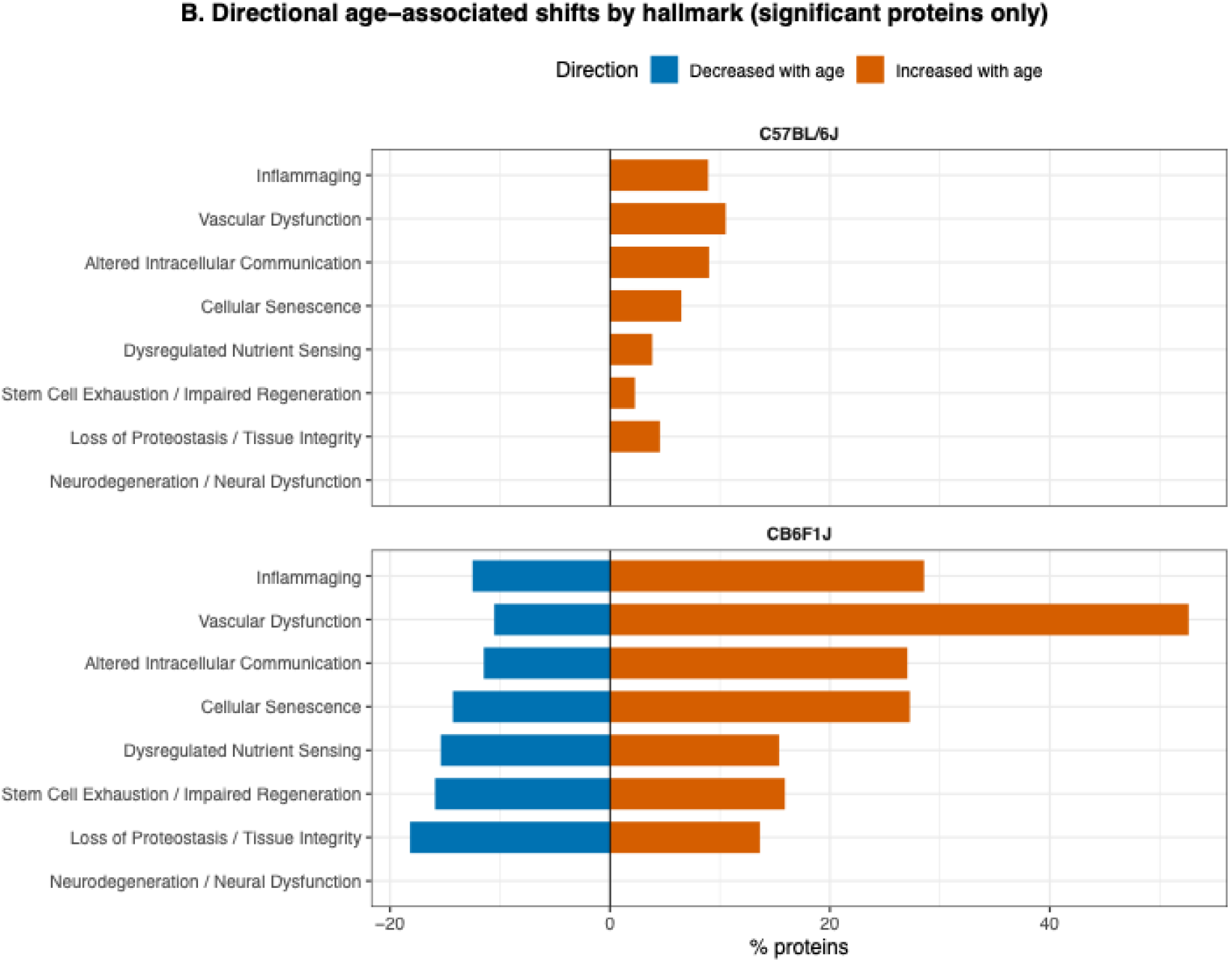
Directional age-associated shifts in circulating proteomic hallmarks across mouse strains (significant proteins only). For each hallmark of aging, the proportion of proteins exhibiting increased (orange) or decreased (blue) expression with age is shown for C57BL/6J (top) and CB6F1J (bottom) mice, restricted to proteins meeting statistical thresholds. Percentages were calculated within each hallmark category. Positive values indicate the fraction of proteins increased with age, whereas negative values indicate the fraction decreased with age. Compared to analyses including all proteins, directional shifts among statistically defined proteins show more selective enrichment across specific pathways, with differences in distribution observed between strains.

**Figure 3.**
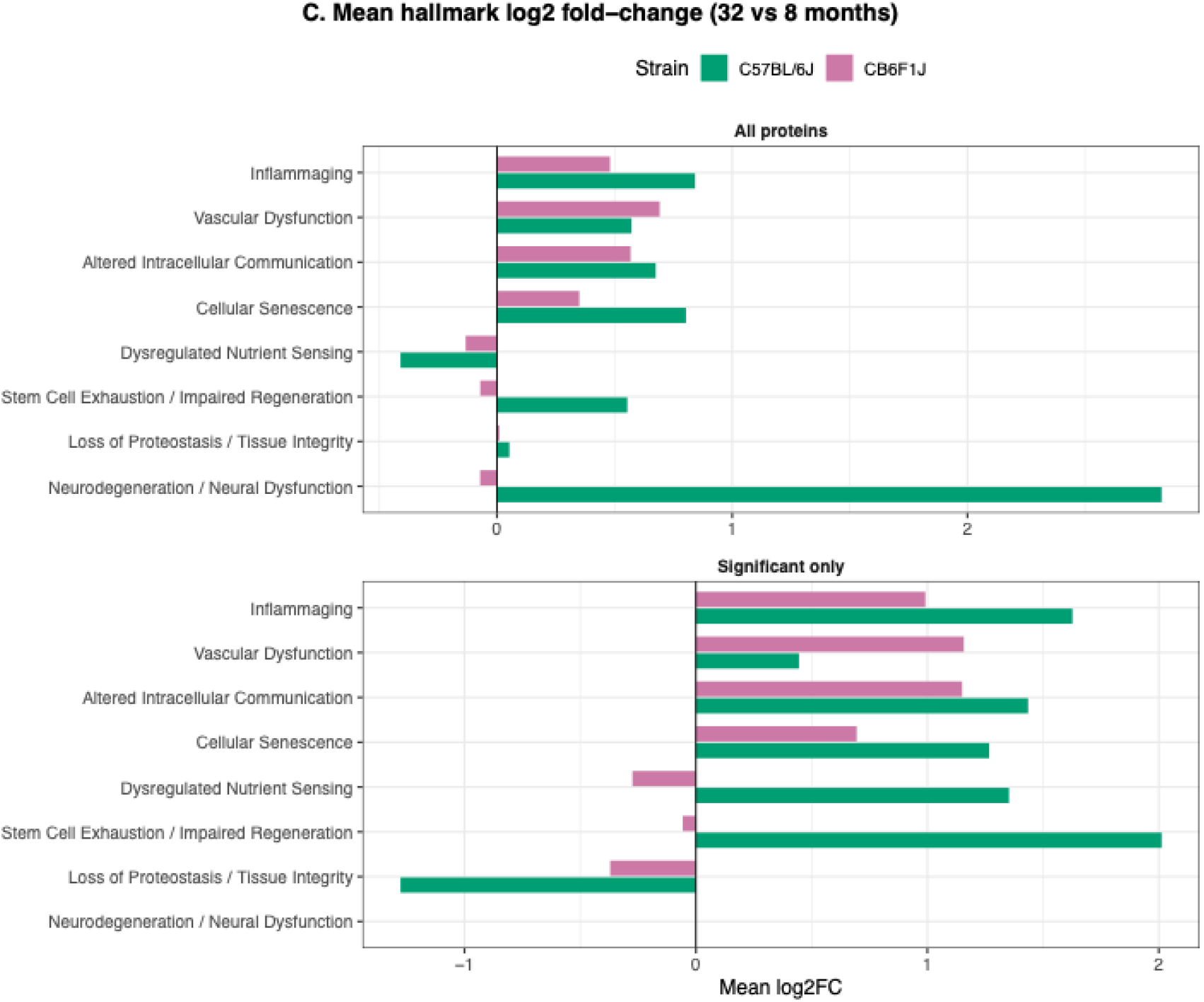
Hallmark-level effect sizes of age-associated proteomic changes across mouse strains. Mean log_2_ fold change (log_2_FC; 32 vs 8 months) is shown for each hallmark of aging in C57BL/6J (green) and CB6F1J (purple) mice. Values are calculated across all proteins (top) and within the subset meeting statistical thresholds (bottom). Positive values indicate higher protein expression with age, whereas negative values indicate reduced expression. Differences in effect size and direction are observed across hallmarks and between strains.

Vascular-associated proteins showed the largest proportion of increases, with 78.9% (15/19) elevated and 52.6% (10/19) represented among the most altered proteins. Corresponding hallmark scores were positive (log_2_FC = 0.69; significant proteins = 1.16). Intercellular communication pathways similarly demonstrated a majority of proteins increased (54.1% [66/122] and 27.1% [33/122]), with positive hallmark scores (log_2_FC = 0.57; significant proteins = 1.15).

In contrast, nutrient sensing pathways showed a predominance of decreased protein levels, with 57.7% (15/26) reduced. Hallmark scores were negative (log_2_FC = −0.13; significant proteins = −0.27). Regenerative pathways also demonstrated a slight negative shift (log_2_FC = −0.07).

Cellular senescence-associated proteins showed 50.6% (39/77) increased, with positive hallmark scores (log_2_FC = 0.35; significant proteins = 0.69). Proteostasis pathways showed minimal net change (log_2_FC = 0.01), with mixed directional changes across individual proteins. Neurodegeneration-associated proteins were limited in number and exhibited variable directional changes.

### C57BL/6J male mice displayed broad directional increases in inflammatory, intercellular, and senescence-associated proteins with heterogeneous changes in metabolic and regenerative pathways

In C57BL/6J male mice, aging was associated with widespread directional shifts across multiple hallmarks (Table 2, Figures 1–3). Inflammatory proteins showed 72.3% (81/112) increased with age, with a smaller proportion represented among the most altered proteins (8.9% [10/112]). Hallmark scores were positive (log_2_FC = 0.84; significant proteins = 1.63).

**Table 2.** Distribution of directional and statistically defined protein changes across hallmarks of aging in C57BL/6J male mice.

| **Hallmark of Aging** | **Total N** | **↑ Proteins (All) (N, %)** | **↑ Proteins (Sig.) (N, %)** | **↓ Proteins (All) (N, %)** | **↓ Proteins (Sig.) (N, %)** |
| --- | --- | --- | --- | --- | --- |
| **Inflammaging** | 112 | 81 (72.32) | 10 (8.93) | 26 (23.21) | 0 (0.00) |
| **Vascular Dysfunction** | 19 | 13 (68.42) | 2 (10.53) | 6 (31.58) | 0 (0.00) |
| **Altered Intracellular Communication** | 122 | 81 (66.39) | 11 (9.02) | 39 (31.97) | 0 (0.00) |
| **Stem Cell Exhaustion / Impaired Regeneration** | 44 | 25 (56.82) | 1 (2.27) | 18 (40.91) | 0 (0.00) |
| **Dysregulated Nutrient Sensing** | 26 | 12 (46.15) | 1 (3.85) | 14 (53.85) | 0 (0.00) |
| **Cellular Senescence** | 77 | 48 (62.34) | 5 (6.49) | 23 (29.87) | 0 (0.00) |
| **Loss of Proteostasis / Tissue Integrity** | 22 | 13 (59.09) | 1 (4.55) | 7 (31.82) | 0 (0.00) |
| **Neurodegeneration / Neural Dysfunction** | 5 | 3 (60.00) | 0 (0.00) | 2 (40.00) | 0 (0.00) |

Vascular-associated proteins showed 68.4% (13/19) increased, with positive hallmark scores (log_2_FC = 0.57; significant proteins = 0.45). Intercellular communication pathways demonstrated 66.4% (81/122) of proteins increased, with positive hallmark scores (log_2_FC = 0.67; significant proteins = 1.44).

Cellular senescence pathways showed 62.3% (48/77) of proteins increased, with positive hallmark scores (log_2_FC = 0.80; significant proteins = 1.27).

Nutrient sensing pathways showed 53.9% (14/26) of proteins decreased overall, with a negative hallmark score across all proteins (log_2_FC = −0.41), while the subset of most altered proteins showed a positive mean shift (1.35). Regenerative pathways showed positive hallmark scores (log_2_FC = 0.55; significant proteins = 2.01).

Proteostasis pathways showed minimal net change across all proteins (log_2_FC = 0.05), with a negative shift among the most altered proteins (−1.28). Neurodegeneration-associated proteins showed variable directional changes and elevated mean log_2_FC (2.83), with limited representation among the most altered proteins.

### Pathway-level directional changes were shared between strains, while protein-level concordance varied across hallmarks

Comparison of CB6F1J and C57BL/6J male mice showed that multiple pathways exhibited similar directional shifts across strains (Table 3, Figure 4). Inflammatory, vascular, and intercellular communication pathways showed positive hallmark scores in both strains.

**Table 3.** Comparison of hallmark-resolved protein shifts and inter-strain concordance in aging CB6F1J and C57BL/6J male mice.

| **Hallmark of Aging** | **Log<sub>2</sub>FC Mean (All)** |  | **Log<sub>2</sub>FC Mean (Sig.)** |  | **Concordance** |  |
| --- | --- | --- | --- | --- | --- | --- |
|  | **CB6F1J** | **C57BL/6J** | **CB6F1J** | **C57BL/6J** | **All** | **Sig.** |
| **Inflammaging** | 0.48 | 0.84 | 0.99 | 1.63 | 0.33 | 0.51 |
| **Vascular Dysfunction** | 0.69 | 0.57 | 1.16 | 0.45 | 0.11 | 0.50 |
| **Altered Intracellular Communication** | 0.57 | 0.67 | 1.15 | 1.44 | 0.24 | 0.31 |
| **Stem Cell Exhaustion / Impaired Regeneration** | -0.07 | 0.55 | -0.06 | 2.01 | -0.10 | 0.00 |
| **Dysregulated Nutrient Sensing** | -0.13 | -0.41 | -0.27 | 1.35 | -0.06 | 0.04 |
| **Cellular Senescence** | 0.35 | 0.80 | 0.69 | 1.27 | 0.38 | 0.69 |
| **Loss of Proteostasis / Tissue Integrity** | 0.01 | 0.05 | -0.37 | -1.28 | 0.41 | 0.77 |
| **Neurodegeneration / Neural Dysfunction** | -0.07 | 2.83 | NA | NA | -0.95 | NA |

**Figure 4.**
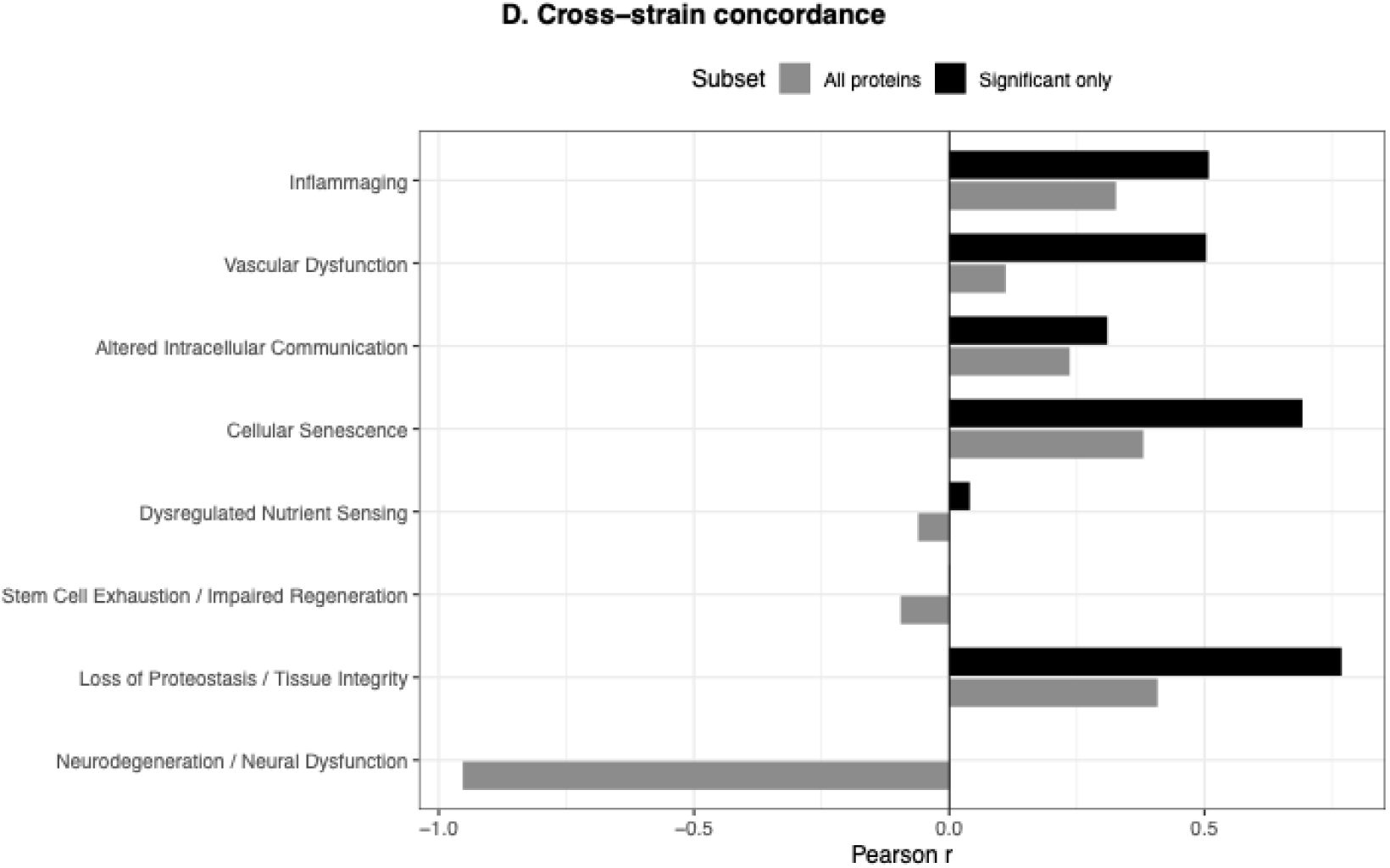
Cross-strain concordance of age-associated proteomic changes across aging hallmarks. Pearson correlation coefficients (r) of protein-level log_2_ fold changes (32 vs 8 months) between CB6F1J and C57BL/6J mice are shown for each hallmark of aging. Concordance was calculated across all proteins (grey) and within the subset meeting statistical thresholds (black). Positive values indicate shared directionality of change across strains, whereas negative values indicate opposing patterns. Concordance varies across hallmarks, with higher agreement observed in select pathways and lower or negative correlations in others.

Protein-level concordance varied across pathways. Correlation coefficients for all proteins were as follows: inflammation (r = 0.33), intercellular communication (r = 0.24), and vascular pathways (r = 0.11). Concordance within the subset of most altered proteins was higher for vascular pathways (r = 0.50).

Nutrient sensing pathways showed low concordance (r = −0.06), and regenerative pathways showed near-zero concordance (r = −0.10), with differing hallmark scores between strains (CB6F1J = −0.07; C57BL/6J = 0.55).

Cellular senescence pathways showed moderate concordance (r = 0.38), with higher concordance among the most altered proteins (r = 0.69). Proteostasis pathways showed low overall change but higher concordance among the most altered proteins (r = 0.77). Neurodegeneration-associated proteins showed variable concordance (r = −0.95), with limited representation.

## Discussion

This study showed that circulating protein array profiles obtained from a single blood draw can resolve hallmark-level features of aging and detect inter-strain heterogeneity, suggesting that aging signatures can be quantified non-invasively within the same organism with repeated assessment across the lifespan.

Aging-associated changes observed in CB6F1J mice were characterized by coordinated increases in inflammatory, vascular, and intercellular signaling proteins, alongside reductions in nutrient sensing and regenerative pathways. These patterns are consistent with established aging biology, including chronic inflammation, vascular remodeling, and altered intercellular communication as central features of organismal aging [2–3]. Prior work has shown that CB6F1J mice exhibit pronounced age-related functional decline, including impairments in cognition, locomotion, and tissue integrity [10–12], alongside transcriptomic signatures of increased inflammatory and degenerative processes that are conserved across mouse strains and tissues with aging [13–14]. The circulating proteome patterns observed here align with these previously reported phenotypes.

In contrast, C57BL/6J mice exhibited broader but more heterogeneous pathway-level changes, including increased inflammatory and senescence-associated proteins with variable regulation of metabolic and regenerative pathways. Previous studies have reported that C57BL/6J mice display relative resilience to certain age-associated pathologies compared to hybrid strains, including slower progression of functional decline and differences in tissue-specific aging trajectories [7]. Although core aging-associated transcriptomic programs, such as inflammatory activation, are shared, transcriptomic and histological analyses demonstrate substantial variation in the magnitude, timing, and pathway composition of aging-related changes across tissues and strains, particularly in metabolic and regenerative pathways [15–16]. The present findings extend these observations by demonstrating that such differences are detectable at the level of circulating proteins.

This is particularly relevant as the use of circulating biomarkers also circumvents the need for terminal tissue collection, which is a major limitation of transcriptomic and histological approaches in animal models [17]. Large-scale efforts such as bulk and single-cell RNA sequencing have provided detailed maps of age-associated changes across tissues, but these methods are inherently invasive and not compatible with longitudinal sampling [14]. In contrast, blood-based profiling has been increasingly recognized as a scalable and translational approach for assessing biological aging, including through proteomic and epigenetic biomarkers [4–5].

Importantly, the ability to resolve hallmark-level changes from circulating proteins suggests that aging processes are reflected in the blood. Previous studies have demonstrated that blood-based proteomes capture age-associated shifts in inflammation, extracellular matrix remodeling, and signaling pathways [5, 18] and can be used to predict biological age and functional decline [19–20]. The present findings extend this framework by demonstrating that such signatures are sufficiently sensitive to distinguish strain-specific aging patterns and detect variability across pathways within individuals.

In summary, these observations support the feasibility of using serum protein profiling as a non-invasive approach to align with biological aging. If translated to human studies, this approach may enable longitudinal assessment of aging trajectories using routine blood sampling.

## Supporting information

Publication License for Graphical Abstract

## Acknowledgments

This work was supported by the National Institute on Aging of the National Institutes of Health under award number R01 057381 (Ladiges, PI). The funder had no role in study design, data collection and analysis, decision to publish, or preparation of the manuscript.

## Availability of Data and Materials

The datasets generated during the current study are available from the corresponding author on reasonable request.

---

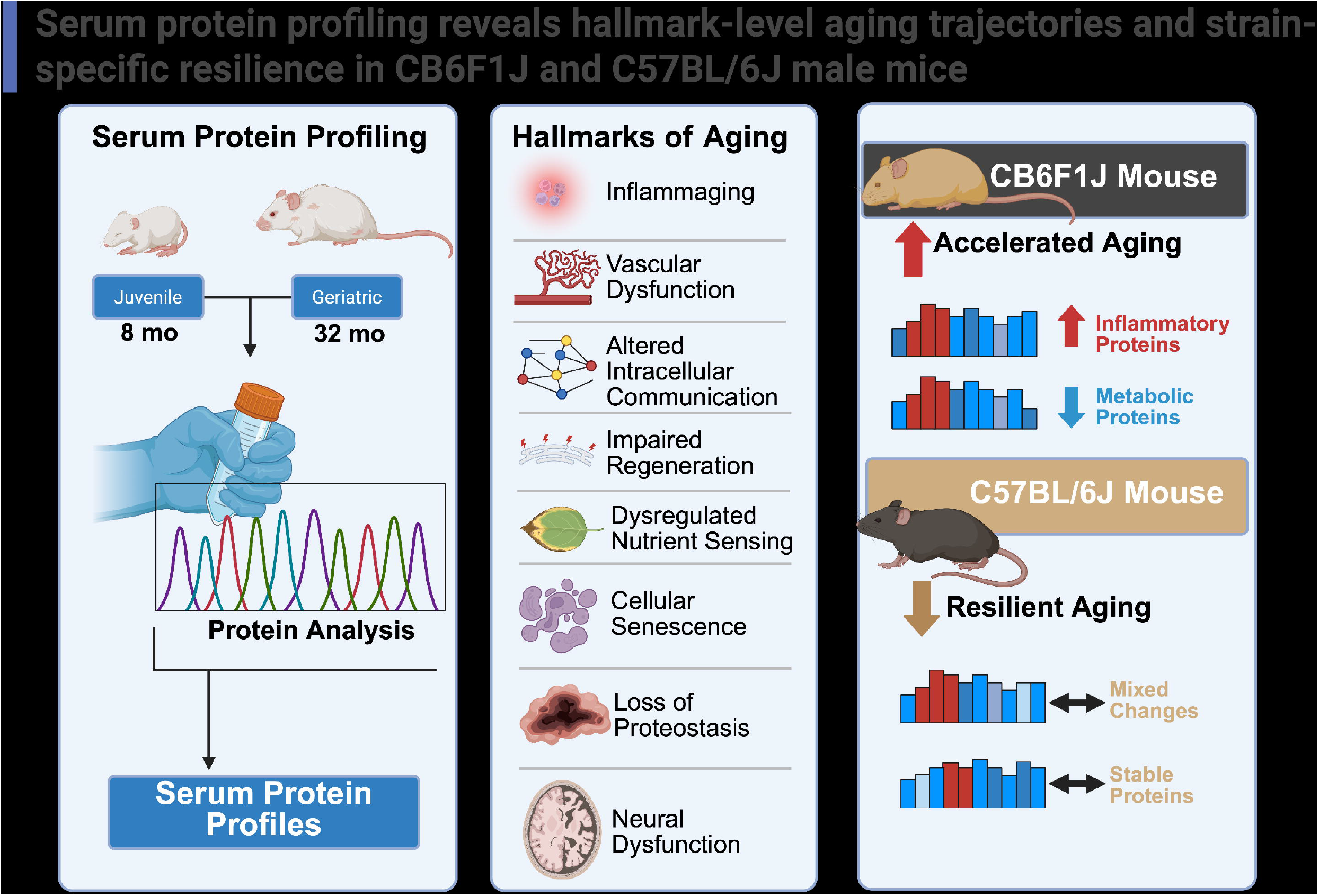

